# Low normal factor V enhances thrombin generation in hemophilia A by a substrate competition mechanism with factor Xa

**DOI:** 10.1101/2022.06.27.496845

**Authors:** Dougald M. Monroe, Christine Baird, Julie A. Peterson, Alan E. Mast, Marilyn Manco-Johnson, Michael Stobb, Suzanne Sindi, Aaron L. Fogelson, Karin Leiderman, Keith B. Neeves

**Affiliations:** UNC Blood Research Center, University of North Carolina at Chapel Hill, Chapel Hill, NC; Hemophilia and Thrombosis Center, University of Colorado Anschutz Medical Campus, University of Colorado Denver | Anschutz Medical Campus, Aurora, CO; Versiti Blood Research Institute, Milwaukee, WI; Department of Cell Biology, Neurobiology and Anatomy, Medical College of Wisconsin, Milwaukee, WI; Department of Pediatrics, Section of Hematology, Oncology, and Bone Marrow Transplant, University of Colorado Denver | Anschutz Medical Campus, Aurora, CO; Department of Mathematics and Computer Science, Coe College, Cedar Rapids, IA; Department of Applied Mathematics, University of California, Merced, Merced, CA; Departments of Mathematics and Biomedical Engineering, University of Utah, Salt Lake City, UT; Department of Applied Mathematics and Statistics, Colorado School of Mines, Golden, CO; Department of Bioengineering, University of Colorado Denver | Anschutz Medical Campus, Aurora, CO

## Abstract

Bleeding patterns in people with hemophilia A cannot be predicted solely by factor VIII (FVIII) levels. Some of the variance in bleeding may be attributed to differences in plasma protein composition, and specifically other coagulation factors where the normal ranges span 50-150% of the population mean. We recently used a mathematical model of thrombus formation that identified factor V (FV) levels as a strong modifier of thrombin generation in FVIII deficiencies. Counterintuitively, the model predicted low normal FV levels enhanced thrombin generation. Here, we tested this prediction and investigated its mechanism. Thrombin generation in plasma from people with FVIII deficiencies (<5%) were negatively correlated with FV levels. A substrate competition mechanism wherein FV and FVIII compete for activation by FXa during the initiation of coagulation was tested in three models: In a purified system containing only FV, FVIII, and FXa, reducing FV enhanced FVIII activation. In synthetic plasma containing the essential proteins of the extrinsic coagulation pathway, low normal FV levels resulted in enhanced thrombin generation both in the presence or absence of TFPIα. In mixture studies using FVIII-deficient human plasma immunodepleted of FV, thrombin generation was enhanced at lower levels of FV. In all models the trend was nonlinear as the effect size was significant at low, but not high, FV levels. Our data show that low normal plasma levels of FV enhance thrombin generation in hemophilia A by reducing FXa substrate competition for FVIII activation and implicate FV levels as a strong modifier of bleeding in hemophilia A.

**Key Points:**

- Low normal levels of FV enhance thrombin generation in hemophilia A by reducing substate competition for FVIII activation.
- Plasma FV levels are a strong modifier of bleeding in hemophilia A.

## Introduction

Hemophilia A is a X-linked recessive genetic disorder that causes a deficiency in coagulation factor VIII (FVIII). The deficiency is clinically defined into three categories; mild (6-30%), moderate (1-5%), and severe (<1%).^1^ Within these categories, FVIII levels are not always accurate predictors of bleeding risk; some individuals with severe deficiencies present with moderate bleeding phenotypes.^2^ Sources of variability within the blood, vascular wall, or extravascular environment may modify the severity of the underlying FVIII deficiency. We previously used a mechanistic mathematical model to examine how variances in coagulation factor levels affect thrombin generation in FVIII deficient blood.^3^ The model results led to the unexpected prediction that FV levels in the lowest quartile of the normal range (50-75%) enhance thrombin generation when FVIII levels are <10%. The objective of this study is to test that prediction and determine the mechanism(s) that explain it.

Our mathematical model of coagulation tracks the concentrations of zymogens, cofactors, complexes, enzymes, and binding sites over time within a small tissue factor (TF) initiated vascular injury under flow.^4,5^ As such, it not only outputs total thrombin generation, but can also suggest the biochemical mechanisms that determine its dynamics. Within the confines of the model, a substrate competition mechanism explains why lower FV levels enhance thrombin generation in hemophilia A: FV and FVIII compete for the initial FXa produced by the TF:FVIIa complex, even before thrombin is generated.^3^ The model shows that FXa is able to convert more FVIII to FVIIIa when lower levels of FV are present. Even though FVIII is deficient in hemophilia A, it is not completely absent, and the formation of more FVIIIa:FIXa following TF-initiated coagulation ultimately enhances the rate and amount of thrombin generation in the mathematical model.

In this study, we tested the hypothesis that FV levels in the low normal range enhance thrombin generation in hemophilia A. We also sought to determine if substrate competition for FXa during coagulation initiation is a feasible mechanism to explain this enhancement in thrombin generation. Consistent with our hypothesis, we find that FV levels are negatively correlated with the peak concentration of thrombin generation within a cohort of people with moderate to severe hemophilia. We then used three biochemical systems—a minimal set of purified proteins, synthetic plasma, and FV immunodepleted from FVIII deficient plasma—to test the feasibility of the FXa substrate competition mechanism. In each of these systems, reducing FV levels below 100% of their normal levels enhanced the velocity and peak height in thrombin generation assays when FVIII levels were ≤ 5%.

## Methods

### Materials

FVIII deficient plasma was purchased from George King Bio-Medical. Factor V antibody for immunodepletion was from Green Mountain Antibodies (GMA-5007). Human factor V ELISA kit was from abcam (ab137976). Dynabead™ coupling kit was from Life Technologies (14311D). Normal pooled plasma was generated in house with the plasma of 38 individuals. Prothrombin, FV, FVII, FX, FXa, FV, FXI, and antithrombin were purchased from Haematologic Technologies (now Prolytix). Recombinant full length TFPIα was expressed in HEK cells. Recombinant human factor IX (Benefix) and factor VIII (Kogenate FS) was from University of North Carolina Pharmacy. Thrombin substrate benzyloxycarbonyl-glycyl-glycyl-arginine 7-amido-4-methylcoumarin (ZGGR-AMC) was purchased from Bachem AG. Phosphatidylcholine, phosphatidylserine, and phosphatidylethanolamine were purchased from Avanti Polar lipids (130601P, 840032P, 850725P) and prepared as previously described.^6,7^ Dade Innovin was purchased from Siemens and used as the TF source in purified and synthetic plasma systems. The FXa substrate Pefachrome FXa was from DSM Nutritional Products (Kaiseraugst, Switzerland).

### FXa substrate competition assays in a purified system

Recombinant FVIII (1 U/mL) was incubated with 50 pM FXa, 4 μM lipid vesicles (Avanti Polar Lipids PS:PC:PE 3:2:5), and FV (0, 6.25, 12.5, 18.75, 25, 31.25, and 37.5 nM). At timed intervals, FVIII activity was assessed by measuring the ability of the FIXa:FVIIIa complex to activate FX (2 nM FIXa, 135 nM FX, 0.33 U/mL FVIIIa) for 30 seconds. The reaction was stopped by addition of an excess of EDTA and the amount of FXa generated was determined by measuring the rate of cleavage of the FXa substrate. The apparent FVIII activation and inactivation were modeled by the following set of ordinary differential equations:

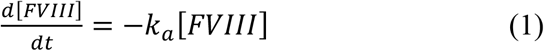

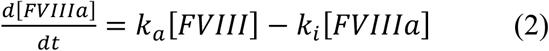

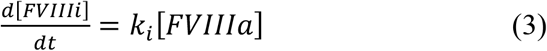

where [FVIII], [FVIIIa], and [FVIIIi] are the concentrations of the zymogen FVIII, activated FVIII, and inactivated FVIII, respectively; *k_a_* and *k_i_* are the activation and inactivation rates. These equations were solved numerically using *lsode* from the FORTRAN library *odepack* in SciPy. The rate constants were estimated by minimizing the sum of squares of differences between the experimental measurements and simulation predictions using the *leastsq* function in SciPy.

### Thrombin generation assays with synthetic plasma

Synthetic plasma contained prothrombin (1.4 μM), FVII (10 nM), FXI (30 nM), FIX (90 nM), FX (140 nM), antithrombin (AT, 3.2 μM), and phospholipids (4 μM, PS:PC:PE 14:41:44), and where noted TFPIα. FVIII concentration was fixed at 0.05 U/mL (5% of the normal plasma level) and FV concentration was varied from 2-12 μg/mL (25%-150% normal plasma level). Zymogen factors were incubated with 10X plasma concentration of AT for at least 2 hours to remove traces of activated factor. Proteins, lipids, TF, and substrate were mixed and thrombin generation was initiated by addition of calcium. Cleavage of the thrombin substrate Z-Gly-Gly-Arg-4-methylcoumaryl-7-amide (416 μM) was measured at 460 nm with excitation at 390 nm in a Synergy H1 plate reader. Data was plotted as the change in fluorescence during each time interval (delta F) and thrombin generation metrics were calculated from custom scripts: The max is the highest value measured for each curve. The lag is the time required for the thrombin concentration to reach 10% of the max value. The velocity is the fit of a straight line through the delta fluorescence values between 10% and 90% of the max.

### Measuring FV, FVIII, and TFPI levels in plasma

FVIII activity was measured using a modified activated partial thromboplastic time assay. Prothrombin and FV activities were measured with modified prothrombin time assays. Both types of assays used calibration curves based on dilutions of normal pooled plasma and prothrombin, FV, and FVIII-deficient plasma using the Compact Max^®^ (Stago) benchtop analyzer following manufacturer’s instructions. These activity assays have a standard deviation of 10%. TFPI and TFPIα were measured by ELISA as described ^8,9^ using mouse monoclonal antibodies directed against the first (MαK1), second (MαK2),^10^ or third (MαK3) Kunitz domains of human TFPIα (all kindly provided by Novo Nordisk, Copenhagen, Denmark). These antigen assays have a standard deviation of 4% and 8%, respectively.

### Factor V immunodepletion

FVIII-deficient plasma (<1% FVIII activity) was mixed with normal pooled plasma to achieve a FVIII activity level of 5% normal. This low FVIII plasma was depleted of FV using a Dynabeads™ coupled to an anti-FV antibody according to manufacturer’s instructions. Following depletion, FV activity was measured with the modified prothrombin time described above. The resulting FV-depleted plasma was mixed with the non-depleted plasma to perform mixture studies.

### Human subjects

Subjects were recruited at the Hemophilia and Thrombosis Center at the University of Colorado Anschutz Medical Campus. Human whole blood was collected into 3.2% sodium citrate. Plasma was isolated from whole blood by centrifugation and stored at −80°C. The study and consent process received from the Colorado Multiple Institutional Review Board in accordance with the Declaration of Helsinki.

### Calibrated automated thrombography

Thrombin generation in plasma was measured with the Calibrated Automated Thrombogram Assay (CAT Assay, Thrombinoscope BV), which consists of a plate reader (Fluroskan Ascent) coupled with the Thrombinoscope software. PPP-reagent (Stago) including 5 pM TF and phospholipid was added to each well followed by plasma and lastly a buffer containing calcium ions. Fluorescence was detected at 390/460 nm ex/em every 20 seconds for 90 minutes. Each condition was run in triplicate.

### Statistical analysis

Pearson correlation coefficients were calculated to measure correlations between plasma coagulation factor levels. All other calculations were done using the SciPy and NumPy packages and plots were generated using Seaborn in Python.

## Results

### FV levels are negatively correlated with thrombin generation metrics in hemophilia A plasma

To test the hypothesis that thrombin generation is inversely correlated with FV levels in hemophilia A, we examined plasmas from a cohort of people with severe to moderate FVIII deficiency (FVIII<5%, n = 75, Table S1). Many of the individuals in this cohort were on prophylactic therapy, so the plasma samples used here were obtained at or near the trough of their dosing regimen. Prothrombin, FV, total TFPI, and TFPIα levels have a normal distribution with means near the normal levels (100%) within this cohort (Fig. 1A, Supplemental Fig. 1). FVIII levels were non-normally distributed with a bias towards the lower end of the range (Supplemental Fig. 1).

**Figure 1.**
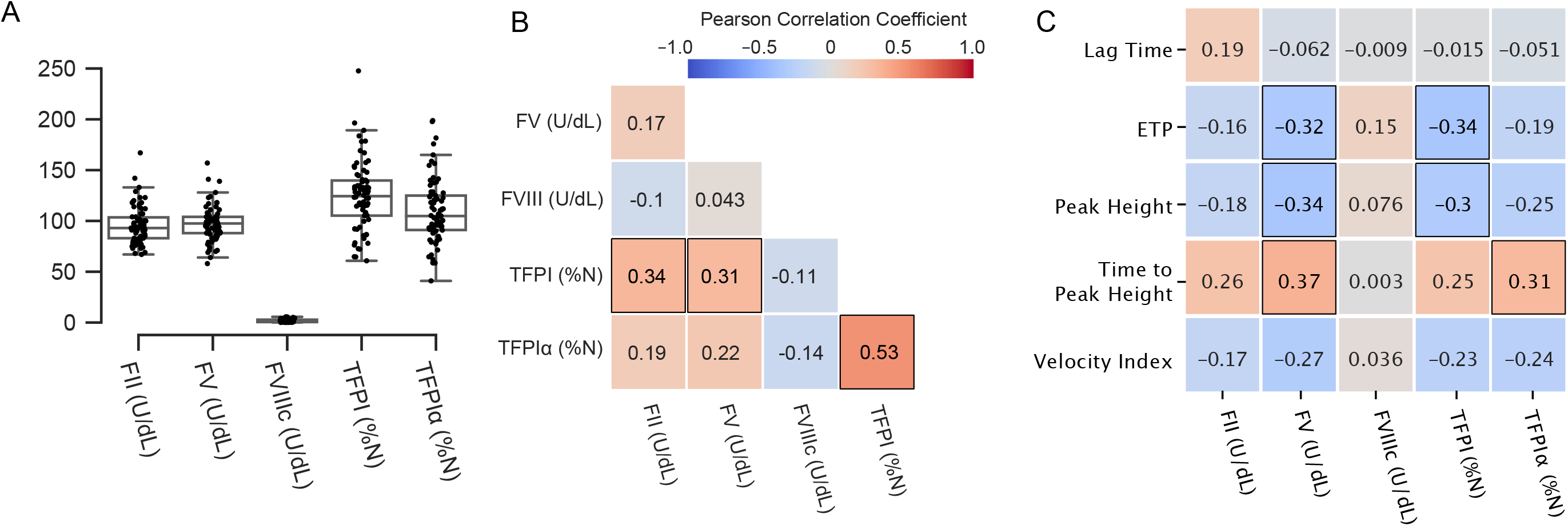
Factor levels in a cohort (n = 75) of individuals with severe and moderate FVIII deficiencies. (A) Distribution of measured factor levels. (B-C) Pearson correlation coefficients between factor levels and factor levels and CAT metrics. See Tables S2 and S3 for p-values for all correlation coefficients. Correlation coefficients with p<0.01 outlined in black.

To determine if these levels (prothrombin, FV, total TFPI, and TFPIα) are related to each other we calculated Pearson correlation coefficients (Fig. 1B, Table S2). There is a significant (p<0.01) positive correlation between total TFPI levels and both prothrombin and FV. There is a weak positive correlation between FV and TFPIα (p = 0.056, Table S2). As expected, TFPI and TFPIα levels are positively correlated (Fig. 1B).

We performed a similar analysis to determine if these protein levels are correlated with thrombin generation metrics. Thrombin generation was measured by CAT using 5 pM TF. Endogenous thrombin potential (ETP) and peak height are negatively correlated with FV and time to peak height is positively correlated with FV (Fig. 1C, Table S3). Samples with FV<100% have significantly higher peak heights (43 nM vs 31 nM, p<0.01) and reach their peaks faster (124 s vs 154 s, p<0.01) than those with FV>100%. Similarly, ETP and peak height are negatively correlated with total TFPI and the time to peak height is positively correlated with TFPIα (Fig. 1C, Table S3). These correlations suggest that the three metrics—ETP, peak height, and time to peak—depend on both FV and TFPI levels. However, because the correlation between FV and TFPI is roughly as strong as their correlation to these three thrombin generation metrics it is challenging to determine if their effects are independent within this data set.

These data demonstrate that FV and TFPI levels are negatively correlated with thrombin generation, although not necessarily independent of each other, in hemophilia A plasma and provide the rationale to further explore potential mechanisms.

### Low FV enhances FVIII activation by FXa in a purified system

One possible mechanism that explains how low FV levels enhance thrombin generation is substrate competition between FV and FVIII for FXa wherein low FV results in higher FVIII activation. To test the feasibility of this mechanism, we measured FVIIIa generation in a purified system containing FV, FVIII, FXa, and lipid vesicles. FV levels were varied from 0-12 μg/mL, which corresponds to 0-150% of normal levels of FV, and FVIII was fixed at 1 U/mL (Fig. 2A). Decreasing levels of FV led to faster generation of FVIIIa (shorter time to peak) and higher peak concentrations of FVIIIa. When these data are fit to an empirical model of activation and inactivation (Eqns. 1-3), we find that the activation rate of FVIII increases roughly 5-fold as FV levels are reduced from 150% (12 μg/mL) to 50% (4 μg/mL) of the normal levels, while the inactivation rate of FVIIIa remains relatively constant over the same range (Fig. 2B). Since the activation of FVIII involves an initial reaction between FXa and FVIII and the initial FVIII levels are not changing, the increased activation rate in our kinetic model is consistent with the idea that there is more FXa available to bind and activate FVIII. These data provide evidence of the biochemical feasibility of the substrate competition mechanism for FXa; decreasing FV levels corresponds to the availability of more FXa for FVIII activation.

**Figure 2.**
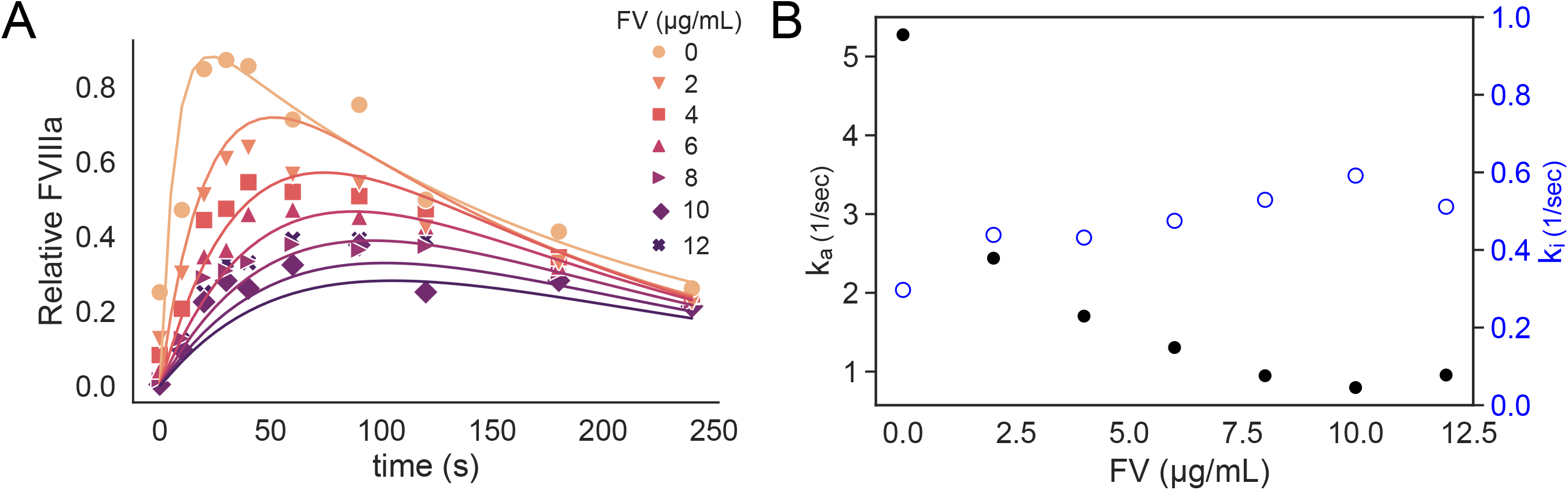
FVIIIa was generated in a purified system of 1 U/mL rFVIII, 50 pM FXa, 0-12 μg/mL FV (0-150% normal levels), and 4 μM phospholipids. (A) At timed intervals FVIII activity was assessed and is reported relative to complete conversion of FVIII to FVIIIa in the purified mixture (symbols). An empirical model of FVIII activation and inactivation was fit to the data to determine rate constants as a function of FV concentration (lines). (B) The rates of activation (ka, black filled circles) and inactivation (ki, blue open circles) of FVIII from fitting to the model described by Eqns. 1-3.

### Low FV enhances thrombin generation in synthetic plasma under hemophilia A conditions

To test whether low to low normal FV concentrations results in significant changes in thrombin generation compared to normal levels of FV, we conducted experiments in a synthetic plasma containing the essential components of the “extrinsic pathway”; prothrombin, TF, FV, FVII, FVIII, FIX, FX, FXI, antithrombin, phospholipids, ± TFPIα. Note that thrombin generation without TFPIα is significantly faster than when it is present, and therefore a lower TF concentration of 1 pM was used in these experiments to observe the initial thrombin dynamics and to yield peak thrombin concentrations comparable to those in the presence of TFPIα at 5 pM TF (Fig. 3A,B).

**Figure 3.**
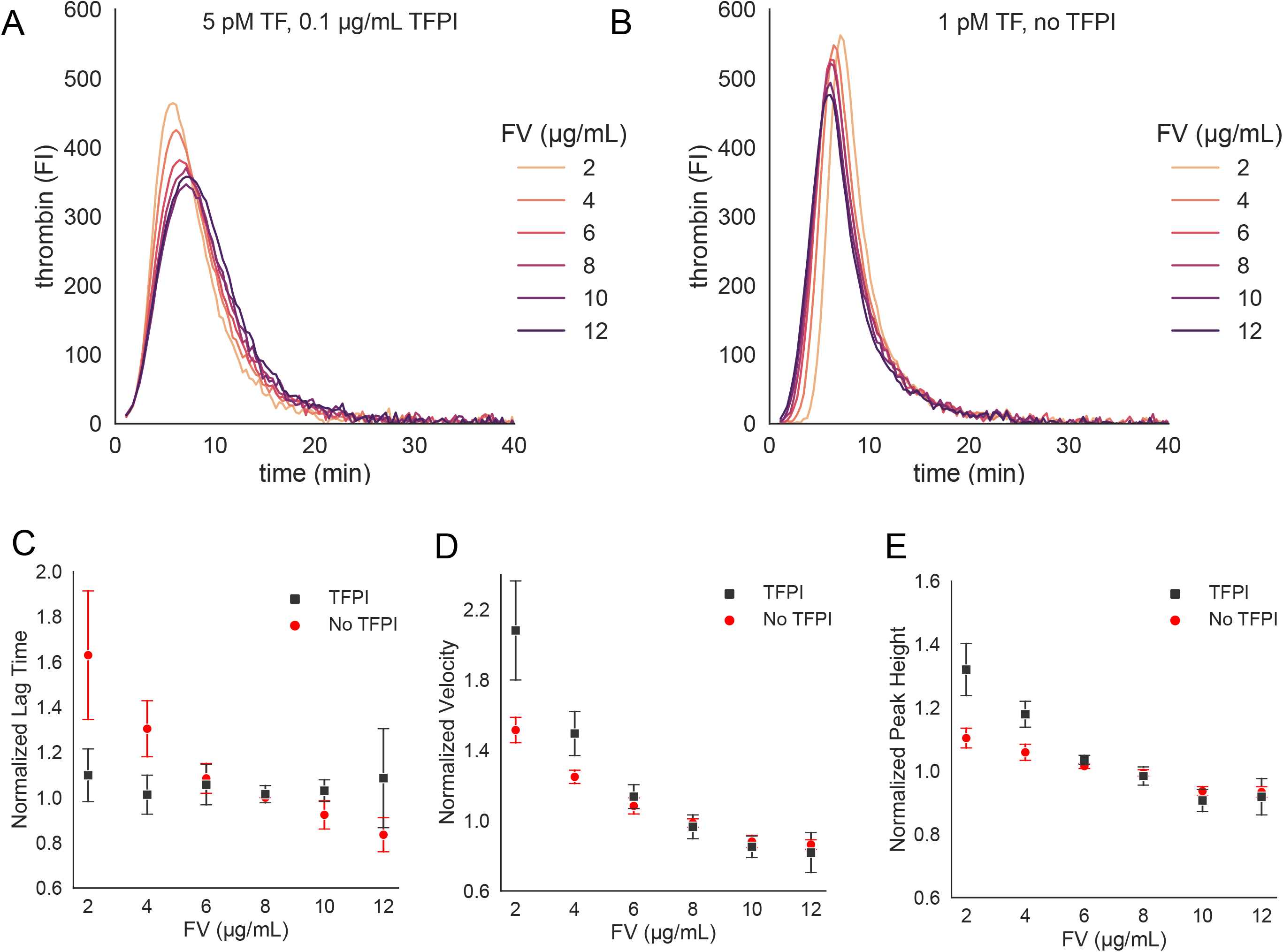
Thrombin generation in synthetic hemophilia plasma (5% FVIII) with varying levels of FV initiated by TF = 5 pM in the presence 0.1 μg/mL TFPIα (A) and initiated by TF = 1 pM in the absence of TFPIα (B). Thrombin generation metrics of lag time (C), velocity (D), and peak height (E) with and without TFPIα. Metrics are normalized by synthetic plasma containing 100% of the population mean concentration of FV (8 μg/mL). Data points represented as mean and standard deviation of n=3.

In the presence of TFPIα, there is an inverse relationship between FV levels and the rate and peak of thrombin generation under hemophilia A conditions (FVIII = 5%), while lag time remains relatively constant under these variations (Fig. 3A,C). There is a 50% increase in velocity, and a 10% increase in the peak height as FV levels were reduced from 8 μg/mL (100%) down to 2 μg/mL (25%) (Fig. 3D,E). Above 8 μg/mL FV, there are little to no additional changes in velocity or peak height.

We next examined whether this relationship between FV and thrombin generation occurs in the absence of TFPIα (Fig. 3B). Similar to experiments with TFPIα, there is an increase in velocity (108%) and an increase in peak height (32%) as FV levels are reduced from 8 μg/mL (100%) down to 2 μg/mL (25%) (Fig. 3D-E). Unlike experiments with TFPIα, there is also a 63% increase in lag time (Fig. 3C), which indicates that reducing FV levels slows initiation.

These findings show that in both the presence and absence of TFPIα, decreasing FV levels enhances thrombin generation under hemophilia A conditions in a synthetic plasma consisting of the extrinsic pathway zymogens, cofactors, and inhibitors and support the substrate competition mechanism between FV and FVIII for FXa, independent of TFPIα.

### Depletion of FV enhances thrombin generation in FVIII deficient human plasma

We next examined whether the trends observed in synthetic plasma are observed in the more complex system of human plasma. Importantly, FVIII is bound to von Willebrand factor (VWF) in plasma which is absent in the synthetic system. FVIII deficient plasma (5% FVIII) containing 80% FV was immunodepleted to yield a FV level of 20%. This low FV/low FVIII plasma (20% FV/5% FVIII) was mixed with normal FV/low FVIII plasma (80% FV/5% FVIII) to yield FV levels of 20%, 40%, 60%, and 80%. In a separate set of experiments, exogeneous FV was added to the normal FV/low FVIII plasma (80% FV, 5% FVIII) to yield FV levels of 100% and 120%. Thrombin generation was measured by CAT for each mixture (Fig. 4A). An inverse relationship between FV levels and lag time, velocity, and peak height was observed in these mixture experiments (Fig. 4B-D). As with the purified and synthetic plasma systems, the relationship between these thrombin generation metrics and FV levels is nonlinear, with the greatest effect size is seen at FV levels below 100%.

**Figure 4.**
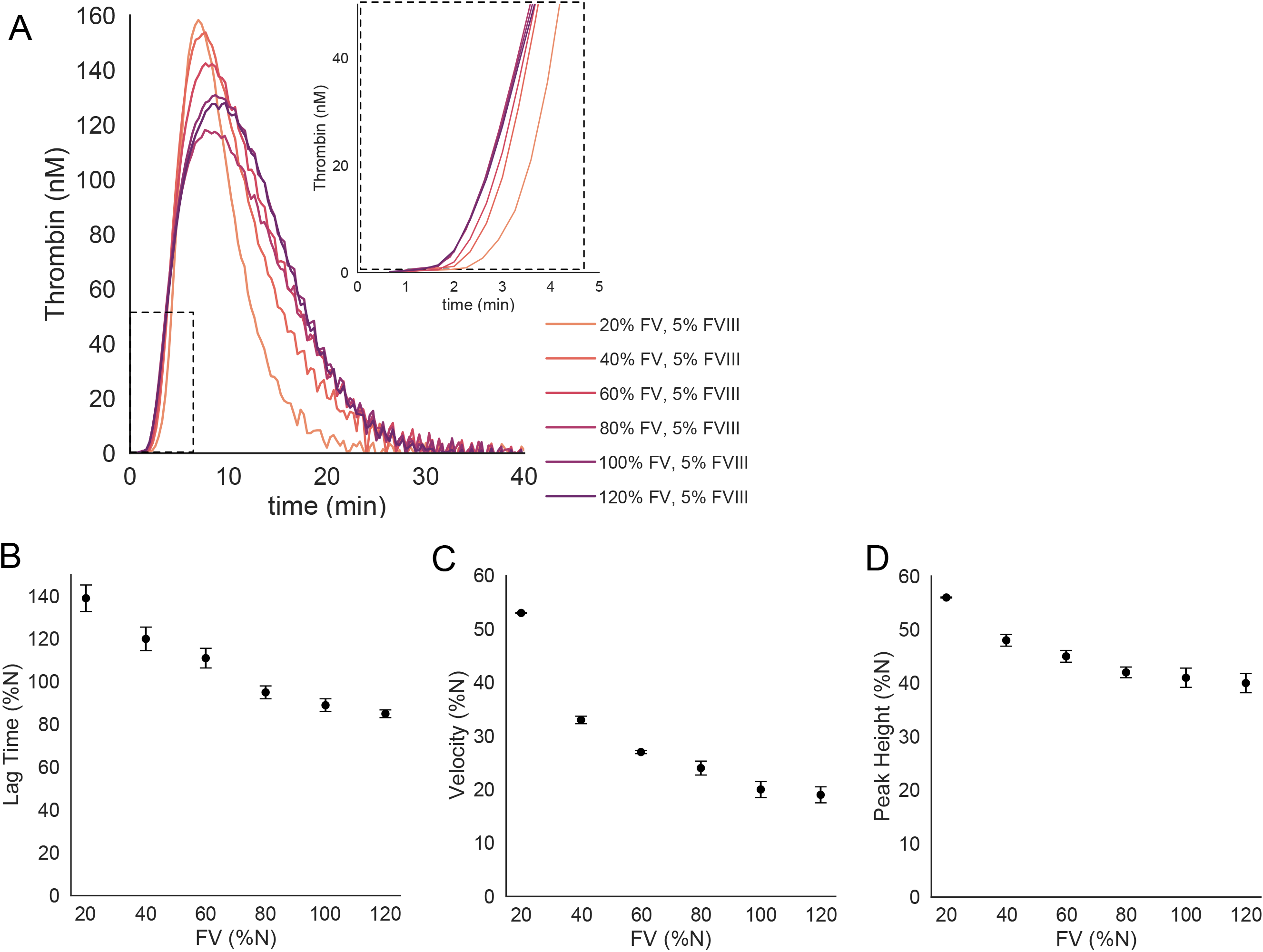
Calibrated automated thrombogram (CAT) with FVIII-deficient (5%) human plasma as function of FV activity. (A) Mixture experiments were conducted with immunodepleted plasma with 20% FV and plasma with 80% FV to yield 40% and 60% FV. Exogenous FV was added to 80% FV plasma to yield 100% FV and 120%. Inset show increased lag time for low FV levels. (B-D) Thrombin generation metrics of lag time, velocity, and peak height relative to normal pooled plasma. Data points represented as mean and standard deviation of n=3.

These data demonstrate that in the complex milieu of human plasma low to low normal FV levels enhance thrombin generation under hemophilia A conditions.

## Discussion

In this study, we provide multiple lines of evidence consistent with a mechanism of substrate competition between FV and FVIII for FXa that drives thrombin enhancement during the initiation of coagulation under hemophilia A conditions. First, in plasma from people with severe to moderate FVIII deficiencies we found that all thrombin generation metrics except time to peak are inversely correlated to FV levels. Second, we show in a purified system that reducing FV enables FXa to generate FVIIIa at a faster rate, consistent with the idea that there is more FXa availability for binding and activating FVIII. Thus, the purified system demonstrates the biochemical feasibility of the substrate competition mechanism using a minimal set of coagulation proteins. Third, in a synthetic plasma system we show that thrombin generation initiated by TF is inversely related to FV levels at low FVIII, both in the presence and absence of TFPIα. Finally, in human plasma with low FVIII we show that immunodepleting FV also results in an inverse relationship between FV levels and thrombin generation metrics.

The substrate competition mechanism highlights the importance of the role of FXa generated by TF:VIIa early in the thrombin generation process (Fig. 5). This FXa activates both FVIII and partially activates FV, the latter of which can bind FXa to form an active prothrombinase complex.^11^ FVIIIa generated in this process can bind FIXa to form the tenase complex, leading to additional FXa formation. With increasing levels of FV, more partially activated FV is produced early and FXa can bind with it, thus moving into prothrombinase complexes and leading to rapid thrombin generation with a short lag phase (Figs. 3B, 4A, 5). With decreasing FV levels, more FXa shifts to activate FVIII, which ultimately leads to more tenase formation and that, in turn, drives greater FXa generation. While this move to FVIII activation leads to a longer lag phase (Figs. 3B, 4A, 5), more prothrombinase complexes are subsequently formed resulting in an increased rate and amount of thrombin generation (Figs. 3C, 4B, 5) even though FV levels are lower. This mechanism may also explain observations that exogenous FV added to combined FVIII and FV deficient plasma reduces peak height in the presence and absence of anti-TFPI antibodies.^12^

**Figure 5.**
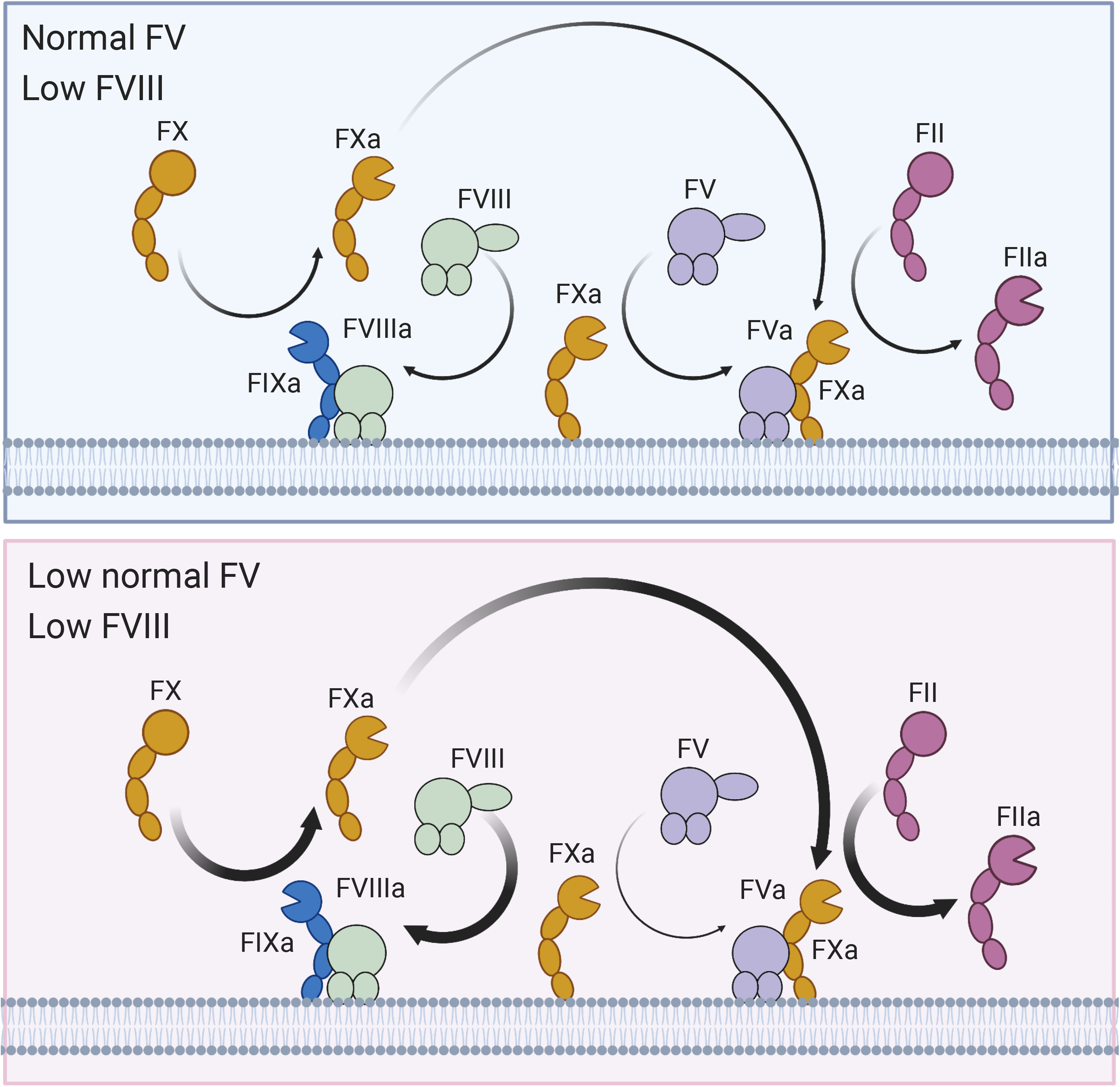
Substrate competition mechanism between FV and FVIII for FXa in hemophilia A. As FV levels are reduced, more FVIII is activated and forms the tenase complex (FVIIIa:FIXa), which increases FXa generation, ultimately leading to more prothrombinase (FVa:FXa) and thrombin generation.

That TFPI can also play an important role is reasonable since it can bind and inhibit FXa and bind to partially activated FV, limiting prothrombinase formation and activity.^13,14^ In synthetic plasma the addition of TFPI led to almost no change in lag times as FV was varied compared to the case with human plasma where lag times were inversely related to FV levels. One explanation for this is that in plasma there is only about 20% of the circulating TFPI that is in its free form,^15^ so the levels of TFPI we used in the synthetic plasma could be much higher than the effective TFPI in the human plasma assay. The higher TFPI in synthetic plasma could be binding most of the FV that is partially activated by FXa early in the process, for all levels of FV, thus rendering the lag times insensitive to changes in FV.

Consistent in both synthetic and FV immunodepleted plasmas was a nonlinear relationship between FV levels and thrombin generation. The effect size was greatest for low to low normal levels of FV and attenuated for FV levels above 100%. This can potentially be explained by relative levels of FV to FVIII in normal plasma compared to hemophilia plasma. In normal plasma, FV levels are much higher compared to FVIII levels, 20-30 nM vs 0.3-0.6 nM, respectively, but the association rates to FXa are similar. In hemophilia, there is roughly a 500-fold difference in these proteins’ levels, thus most of the FXa is occupied by FV. Reducing FV levels frees up FXa for FVIII binding, however increasing it does not have a significant effect on the already low conversion rate of FVIII to FVIIIa. FV and FVIII are not the only binding partners for FXa, TFPI can also bind to and inhibit FXa. Our data in the synthetic plasma shows that the addition of TFPI increases the lag time for thrombin generation as FV levels drop, which could indicate that more FXa is inhibited by TFPI.

The determinants of thrombin generation are multifactorial and, in this study, we have focused on a few proteins (FV and TFPI) to test our overall hypothesis about the correlations seen in patient plasma. The cohort is heterogenous with respect to age (2-75 years old) and it is known that the expression of coagulation proteins varies across the human lifespan.^16,17^ The cohort were also receiving many different replacement products for both on demand and prophylactic therapy. While there were only a few samples with low normal FV levels (50-75%), the correlation between FV and thrombin generation are significant and comparable to other reports. For example, Chelle et al. examined the correlation between thrombin generation and coagulation factor levels in a cohort of people with hemophilia A (n=40).^18^ They found that TFPIα, was the strongest determinant of ETP and peak thrombin in thrombin generation assays initiated by 1 pM TF, with FV levels being the second strongest determinant of peak thrombin. All other coagulation factors had statistically insignificant correlations with thrombin generation metrics in their study. Our results are in general agreement with Chelle et al. as we found both FV and TFPI negatively correlated with thrombin generation initiated by 5 pM TF.

TFPIα binds to FV-short, an alternatively spliced form of FV with an estimated circulating concentration of 0.2 nM.^19,20^ It acts in synergy with protein S to enhance TFPIα activity.^21,22^ Duckers et al. reported a strong correlation between FV and TFPIα across individuals with severe FV deficiencies, partial FV deficiencies, and normal FV levels (n=47, r=0.79).^23^ The strength of this correlation is anchored by data from those individuals with severe FV deficiencies, wherein FV-short levels may also be very low. In the Leiden Thrombophilia Study, FV levels were correlated with TFPIα (r=0.39) in healthy controls (n=474).^24^ Ellery et al. reported a weaker correlation (r=0.18) between the two proteins in healthy subjects (n=435). Here, we found in a cohort of individuals with severe to moderate FVIII deficiencies (n=75) a correlation between FV and TFPIα (r = 0.22) in line with Ellery et al. among individuals with FV levels within the normal range. ^25^

This study focused on the biochemical aspects of coagulation with an emphasis on thrombin generation. Our approach does not account for the role of blood cells, and platelets in particular, on thrombin generation. Platelet α granules contain a partially activated form of FV that contributes to thrombin generation.^26^ Plasma FV and platelet FV levels are correlated with each other,^25^ so it is reasonable to speculate that the trends observed in this study in plasma would translate to a system with platelets. We have previously reported that inhibition of FV in whole blood flow assays enhances fibrin deposition on collagen-TF surfaces, although the relative contributions of plasma and platelet FV are not determined in this assay.^3^ Further investigations are needed to determine how platelet FV contributes to thrombin generation in hemophilia A under varying plasma FV levels. TFPIα is also sequestered and released by activated platelets, and this platelet TFPIα modifies bleeding in FVIII deficient mice.^27^ The role of FV as a cofactor for activated protein C (APC) is another area for future exploration,^28^ as reduced FV levels could also enhance FVIIIa:FIXa formation by reducing FVIIIa degradation via APC; although since thrombin production is a precursor for APC formation, this effect would likely have little influence on early FVIIIa:FIXa formation.

In summary, we have examined how FV levels modulate thrombin generation in the context of FVIII deficiencies. A substrate competition mechanism between FV and FVIII for FXa appears to be biochemically feasible and consistent with data collected in four different experimental systems. The data also supports mechanisms related to TFPIα inhibition of extrinsic tenase and prothrombinase as an important regulator of thrombin generation in FVIII deficiencies.

## Acknowledgments

This study was supported by research funding by the National Institutes of Health grants R01HL151984 and R33HL141794 (A.L.F., K.L., and K.B.N.).

## Authorship

C.B. and J.A.P. performed experiments and analyzed data; A.E.M., M.M.J., M.S., and S.S. contributed to design and analysis of experiments; A.L.F. and K.L. conceived of the research and interpreted data; D.M.M. and K.B.N. conceived of the research, performed experiments, analyzed data, and wrote the manuscript; and all authors contributed to writing the final version of the manuscript.

## Conflict-of-interest disclosure

The authors declare no competing financial interests.

## Notes

### Competing Interest Statement

The authors have declared no competing interest.

