## Supplemental Materials for "Low normal factor V enhances thrombin generation in hemophilia A by a substrate competition mechanism with factor Xa"

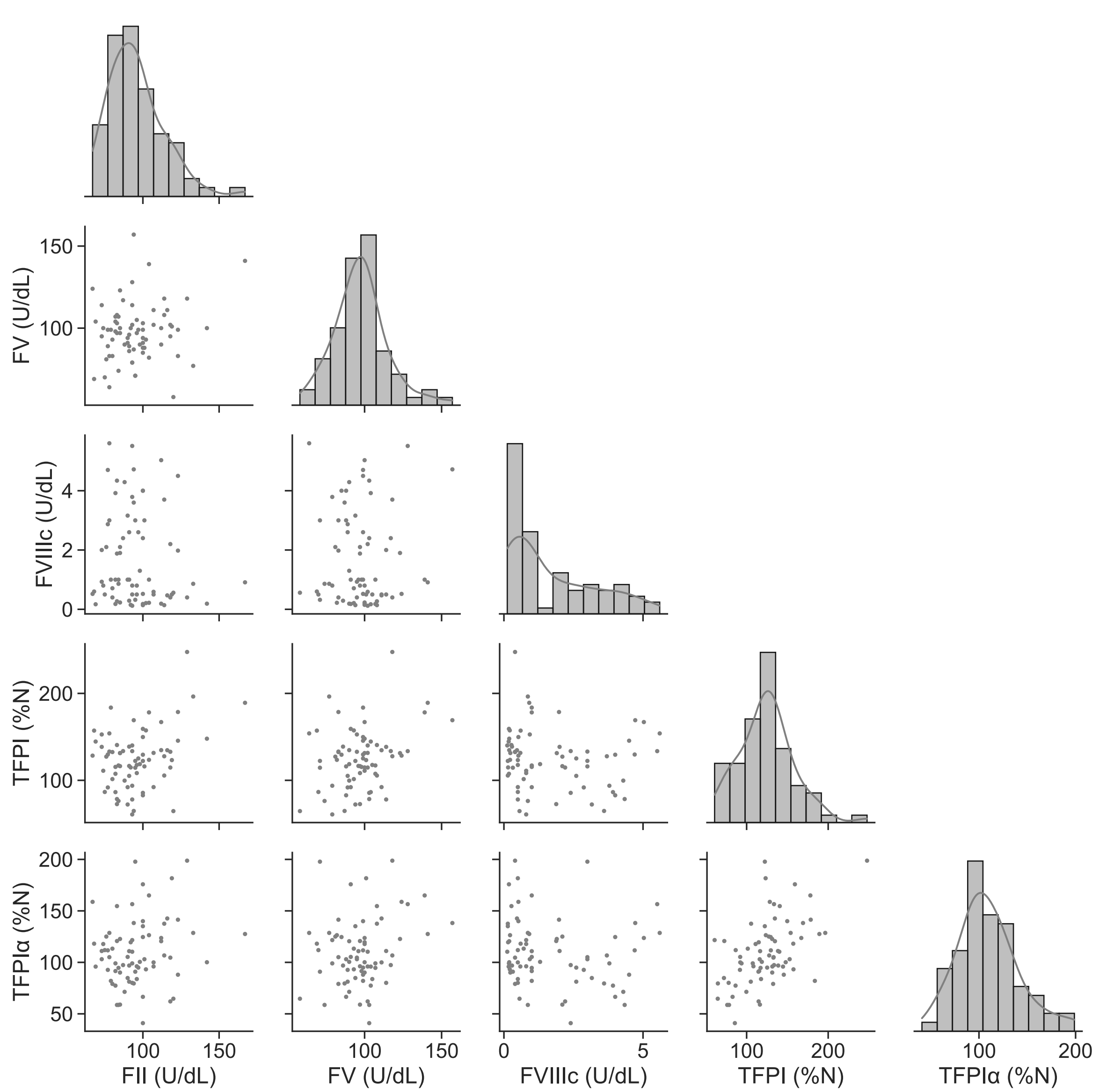


**Supplemental Figure 1.** Histograms of factor level distributions with overlaid kernel density estimation of the probability density function along the diagonal and bivariate plots for pairs of factors in the lower triangle.

**Table S1.** Characteristics of study population (n=75). Data presented as average ± standard deviation and with range in parentheses.

| | **Mean (SD) [Min-Max]** |
|---|---|
| Age | 22.1 (17.6) [2-75.2] |
| BMI | 22.3 (6.0) [13.9-41.3] |
| Prothrombin (U/dL) | 95.5 (18.2) [67-167] |
| FV (U/dL) | 98.0 (17.5) [58-157] |
| FVIII (U/dL) | 1.7 (1.6) [0.1-5] |
| TFPI (% normal) | 125 (33.9) [60.7-247] |
| TFPIα (% normal) | 109 (31.8) [41.0-199] |

**Table S2.** Pearson correlation coefficients between factor level pairs and corresponding p-values for hemophilia A cohort.

| Pair | Correlation Coefficient | P - value |
|---|---|---|
| TFPIα-TFPI | 0.53 | $1.0 \times 10^{-6}$ |
| TFPI-FII | 0.34 | $3.3 \times 10^{-3}$ |
| TFPI-FV | 0.31 | $6.5 \times 10^{-3}$ |
| TFPIα-FV | 0.22 | 0.056 |
| FII-FV | 0.17 | 0.14 |
| TFPIα-FII | 0.19 | 0.11 |
| FVIII-FV | 0.043 | 0.71 |
| FVIII-FII | -0.10 | 0.39 |
| FVIII-TFPI | -0.11 | 0.35 |
| FVIII- TFPIα | -0.14 | 0.25 |

**Table S3.** Pearson correlation coefficients between 5 pM TF CAT metrics and factor levels hemophilia A cohort. Correlation coefficient (p-value).

| | **Prothrombin** | **FV** | **FVIII** | **TFPI** | **TFPIα** |
|---|---|---|---|---|---|
| Lag time | 0.19 (0.11) | -0.062 (0.59) | $9.1 \times 10^{-3}$ (0.93) | -0.015 (0.90) | -0.050 (0.67) |
| ETP | -0.16 (0.17) | 0.32 ($4.9 \times 10^{-3}$) | 0.14 (0.21) | -0.34 ($3.4 \times 10^{-3}$) | -0.19 (0.10) |
| Peak Height | -0.17 (0.13) | -0.34 ($2.6 \times 10^{-3}$) | 0.076 (0.52) | -0.29 (0.010) | -0.25 (0.035) |
| Time to Peak Height | 0.26 (0.02) | 0.37 ($1.0 \times 10^{-3}$) | $3.0 \times 10^{-3}$ (0.98) | 0.25 (0.03) | 0.31 ($6.3 \times 10^{-3}$) |
| Velocity Index | -0.17 (0.13) | -0.27 (0.017) | 0.04 (0.76) | -0.23 (0.047) | -0.24 (0.038) |
